# Characterisation of gene expression markers and glucosinolates during discrete infection stages of *Pyrenopeziza brassicae* in *Brassica napus*

**DOI:** 10.64898/2026.07.15.736298

**Authors:** Ajisa Muthayil Ali, Laura Gimenez Molina, Christoph Crocoll, Aiming Qi, Barbara A. Halkier, Henrik U. Stotz, Rachel Wells

## Abstract

Light leaf spot (LLS), caused by subcuticular hemibiotrophic ascomycete fungus *Pyrenopeziza brassicae*, is a major constraint on oilseed rape (*Brassica napus*) production, yet the genetic and biochemical mechanisms of quantitative disease resistance (QDR) remain poorly defined. Here, disease phenotyping, pathogen quantification, microscopy, gene expression profiling and glucosinolate (GSL) analysis were integrated to dissect resistance mechanisms in *B. napus*. Disease assays of 19 diverse lines revealed clear contrasts between susceptible and resistant genotypes, with the commercial cultivar Ambassador showing a phenotype inconsistent with the UK Recommended List rating. Microscopy demonstrated that resistance within doubled haploid line Cubs Root does not inhibit spore germination or penetration but restricts hyphal branching and subcuticular colonisation from 4 to 8 days post-inoculation. Expression profiling of seven candidate gene expression markers (GEMs) and *pathogenesis-related PR1* showed that *cinnamate-4-hydroxylase*, *phospholipase C4*, *β-adaptin*, *universal stress protein* and the *40S ribosomal subunit protein S24* were strongly pathogen-induced in resistant lines, whereas a *BAHD acyltransferase*, a putative susceptibility factor, was induced only in susceptible cultivars. GSL profiling identified negative correlations between disease severity and total GSLs, particularly aliphatic and aromatic GSLs, with 2□phenylethyl and 7-methylsulfinyl heptyl GSLs showing the strongest associations with resistance. Together, these results highlight coordinated transcriptional and metabolic responses that limit pathogen proliferation and provide targets for breeding durable LLS resistance in *B. napus*.

## Introduction

Oilseed rape (OSR, *Brassica napus*) is a globally important crop used for edible and industrial oil, biodiesel, and animal feed (Karandeni Dewage *et al*., 2018a). However, productivity is constrained by susceptibility to biotic stresses that can cause yield losses of 20–30% (Akter *et al*., 2025). Light leaf spot (LLS), caused by *Pyrenopeziza brassicae*, is a major yield-limiting disease in northern Europe, responsible for losses of up to £160 million annually (Ashby, 1997; Karandeni Dewage *et al*., 2018; Zheng et al., 2020). In the UK, OSR cultivars are evaluated annually under the AHDB (Agriculture and Horticulture Development Board) Recommended List (RL), which provides disease resistance ratings including a 1–9 light leaf spot resistance score to guide growers’ variety choice (AHDB Cereals & Oilseeds, 2026).

*Pyrenopeziza brassicae* is a heterothallic pathogen, requiring two compatible mating types (MAT1 and MAT2) for sexual reproduction, and survives between growing seasons on infected crop debris as pseudothecia. These pseudothecia release wind-dispersed ascospores that initiate primary infections, typically during autumn following crop establishment, by infecting leaves directly through the cuticle. Following penetration, subcuticular proliferation occurs, leading to lesion formation. During the growing season, acervuli form on lesions and produce conidia that are dispersed by rain splash within the same plant or onto neighbouring plants, resulting in secondary infection cycles (Gilles *et al*., 2001; Boys *et al*., 2007). Key pathogenicity factors include the cutinase, Pbc1, for initial penetration (Li *et al*., 2003), the protease, Psp1, expressed during subcuticular colonization (Batish et al., 2003), and cytokinin production, which alters host metabolism to promote pathogen growth (Ashby, 1997; Ashby, 2000).

Quantitative disease resistance (QDR) is generally considered a more durable and non-race-specific form of resistance because it restricts pathogen growth and reproduction rather than completely preventing infection, in contrast to major *R* gene mediated resistance that can be rapidly overcome by pathogen evolution (Corwin and Kliebenstein, 2017). Consequently, pathogen adaptation to QDR is generally slower and less likely to result in complete resistance breakdown (McDonald and Linde, 2002). LLS is a polycyclic disease for which no major *R* genes have been identified, but several loci contributing to quantitative resistance have been mapped (Fell *et al*., 2023).

Evidence indicates that QDR operates at multiple stages of infection. Although *P. brassicae* can infect and colonise both resistant and susceptible hosts, resistant lines support lower pathogen biomass accumulation and reduced sporulation, suggesting that defence primarily limits pathogen colonisation and reproductive development rather than preventing infection (Karandeni Dewage *et al*., 2022). Similar patterns have been reported for the subcuticular pathogen *Rhynchosporium commune*, where resistance in barley (*Hordeum vulgare*) typically restricts fungal growth and sporulation following penetration rather than blocking pathogen entry (Avrova & Knogge, 2012; Zhan *et al*., 2008). Furthermore, distinct loci may control different components of resistance that influence pathogen reproduction or host necrotic responses (Karandeni Dewage *et al*., 2022).

These traits are typically quantitative and polygenic, with loci often mapped to QTL, each contributing partial restriction of pathogen growth, subcuticular proliferation or lesion expansion (Boys *et al*., 2007; Fell *et al*., 2023). Combining multiple QTL can produce strong resistance, although environmental effects and epistatic interactions influence expression (Niks *et al*., 2015; Calenge *et al*., 2005). Mechanistically, QDR is frequently associated with basal defence pathways such as PAMP-triggered immunity involving receptor-like kinases, pathogenesis-related proteins, and hormone-regulated defence signalling such as salicylic acid and ethylene (Boyd *et al*., 2013; Corwin *et al*., 2017).

Recent studies using high-throughput phenotypic screening of 195 *B. napus* lines identified four QDR loci and eight gene expression markers (GEMs) significantly correlated with lower LLS disease severity or higher susceptibility (Fell et al., 2023). Among the eight GEMs, the expression levels of seven were positively correlated with QDR. These genes included *phospholipase C4* (*PLC4*), *cinnamate-4-hydroxylase* (*C4H*), *SNF1 kinase homolog 10* (*KIN10*) and other genes involved in signalling, metabolism, vesicle trafficking, translation or stress responses (Fell *et al*., 2023). One gene, encoding an acyltransferase belonging to the <u>B</u>enzyl alcohol O-acyltransferase, <u>A</u>nthocyanin O-hydroxycinnamoyltransferase, N-<u>H</u>ydroxycinnamoyl/benzoyltransferase and <u>D</u>eacetylvindoline 4-O-acetyltransferase (BAHD) family of proteins (Ma *et al*., 2005), was identified as a potential susceptibility factor (Fell *et al*., 2023).

Glucosinolates (GSLs) are sulfur-containing secondary metabolites characteristic of Brassicaceae that play key roles in plant defence through the production of bioactive hydrolysis products. GSLs and their hydrolysis products may confer biochemical resistance against *P. brassicae*, as they inhibit fungal growth and may act synergistically with QDR genes (Giamoustaris & Mithen, 1997; Stotz *et al*., 2011a; Rahmanpour *et al*., 2009). The interactions and cross-talk between QDR genes, such as *BAHD acyltransferase*, *C4H*, and *PR1*, indicate a complex regulatory network linking defence signalling, metabolite production and pathogen inhibition (Kim *et al*., 2015; Kim *et al*., 2020; Becker *et al*., 2017).

Specifically, we asked (i) whether previously identified QDR-associated GEMs, the *BAHD acyltransferase* and the salicylic acid marker *PR1* are differentially regulated across discrete stages of *P. brassicae* infection in contrasting *B. napus* lines, (ii) at which infection stage resistance is expressed at the cellular level, and (iii) whether glucosinolate profiles are associated with disease phenotype and could reinforce QDR. To address these questions, we phenotyped 19 diverse *B. napus* lines including the twelve accessions we characterised previously (Fell *et al*., 2023), four modern commercial cultivars and three transformable lines, and selected a contrasting subset of eight lines spanning susceptible, intermediate and resistant phenotypes for in-depth pathogen quantification, microscopy, expression profiling of eight markers across four infection stages, and glucosinolate analysis. This integrated design directly tests whether our previously nominated GEMs function as *in planta* resistance markers and link them to a defence-metabolite framework that could guide marker-assisted breeding for durable LLS resistance. An overview of the experimental design and the *B. napus* lines used in each module is shown in Figure 1.

**Figure 1.**
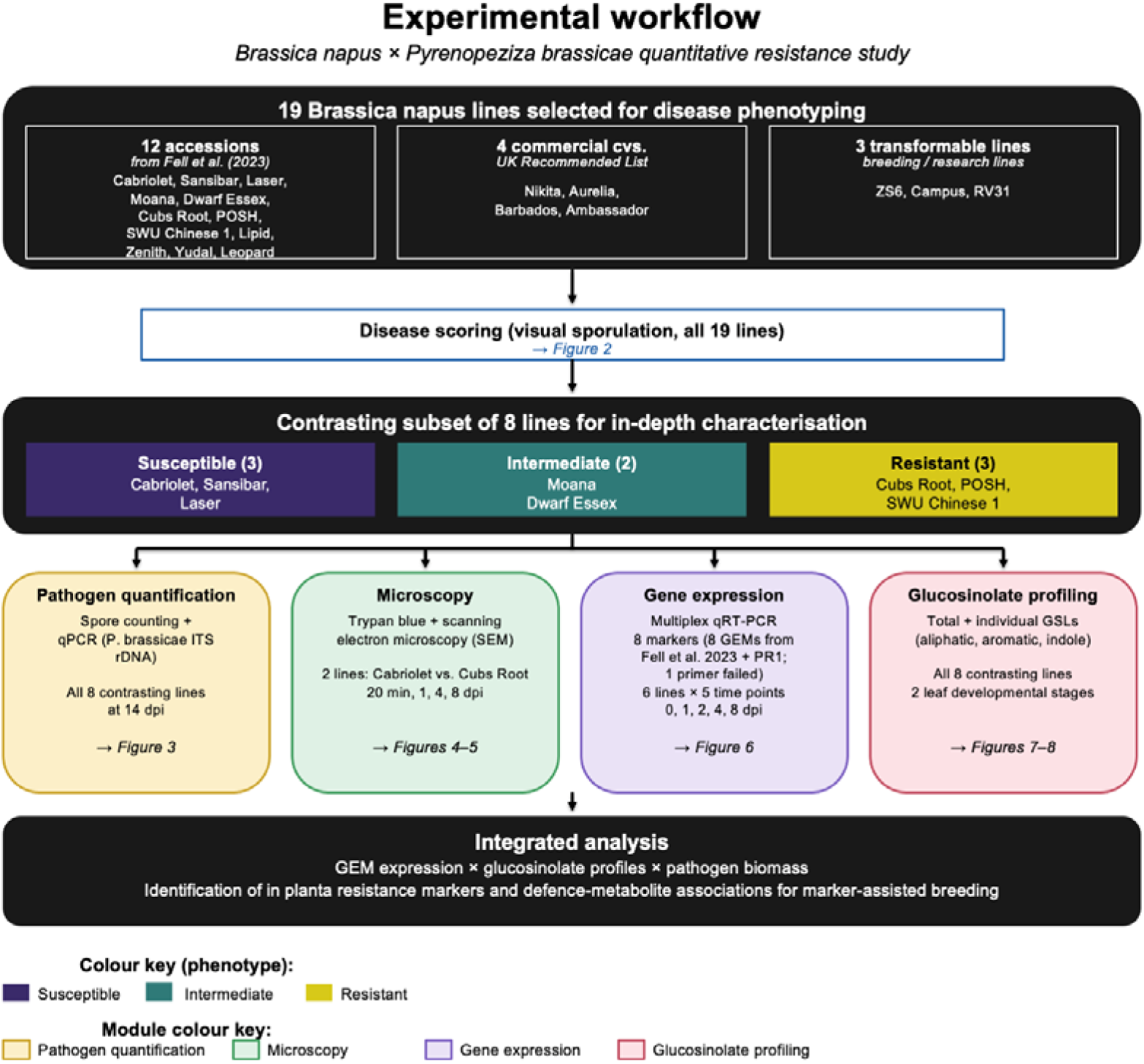
Schematic overview of the experimental design and *Brassica napus* lines used in each module of this study. Nineteen *B. napus* lines were selected for disease phenotyping: twelve from Fell et al. (2023) (Cabriolet, Sansibar, Laser, Moana, Dwarf Essex, Cubs Root, POSH, SWU Chinese1, Lipid, Zenith, Yudal and Leopard), four modern UK commercial cultivars (Nikita, Aurelia, Barbados and Ambassador), and three transformable lines (ZS6, Campus and RV31). All 19 lines were scored for disease severity under controlled-environment conditions (Figure 2). A subset of eight lines spanning the disease spectrum (Cabriolet, Sansibar and Laser [susceptible]; Moana and Dwarf Essex [intermediate]; Cubs Root, POSH and SWU Chinese1 [resistant]) was selected for four experimental modules: (i) pathogen quantification by spore counting and qPCR of *P. brassicae* ITS rDNA at 14 dpi (Figure 3); (ii) microscopy of fungal development by trypan blue staining and scanning electron microscopy in Cabriolet vs. Cubs Root at 20 min, 1, 4 and 8 dpi (Figures 4–5); (iii) multiplex qRT-PCR profiling of eight gene expression markers (GEMs; Fell et al., 2023; *KIN10* excluded due to primer failure) in six lines at 0, 1, 2, 4 and 8 dpi (Figure 6); and (iv) glucosinolate profiling of aliphatic, aromatic and indole glucosinolates in all eight lines across two leaf developmental stages (Figures 7–8).

## Results

### Controlled disease assays identify contrasting *B. napus* lines for studying the interaction with *P. brassicae*

Nineteen *B. napus* lines were selected for disease scoring to determine their susceptibility to *P. brassicae* (Figure 1; Supplementary Table 1). Assessment of the percentage of visual sporulation on infected leaves (Fell *et al*., 2023) classified each line into a susceptible, intermediate or resistant category when using glasshouse conditions for plant growth (Figure 2A). This classification was confirmed when using spore counting as an alternative, more quantitative disease assessment method (Guimaraes *et al*., 2004; Figure 2B). It was noted that cv. Ambassador, despite having high resistance scores in the UK AHDB Recommended Variety List (AHDB Cereals & Oilseeds, 2016), was highly susceptible to *P. brassicae*, whereas cvs. Nikita, Aurelia, and Barbados were consistently resistant. The results obtained under glasshouse conditions were confirmed in a field trial, showing that leaves of cv. Ambassador were more susceptible to *P. brassicae* than the resistant cvs. Nikita, Barbados and Aurelia (Supplementary Figure 1). This indicates effective resistance has not been consistently selected for in modern breeding material.

**Figure 2.**
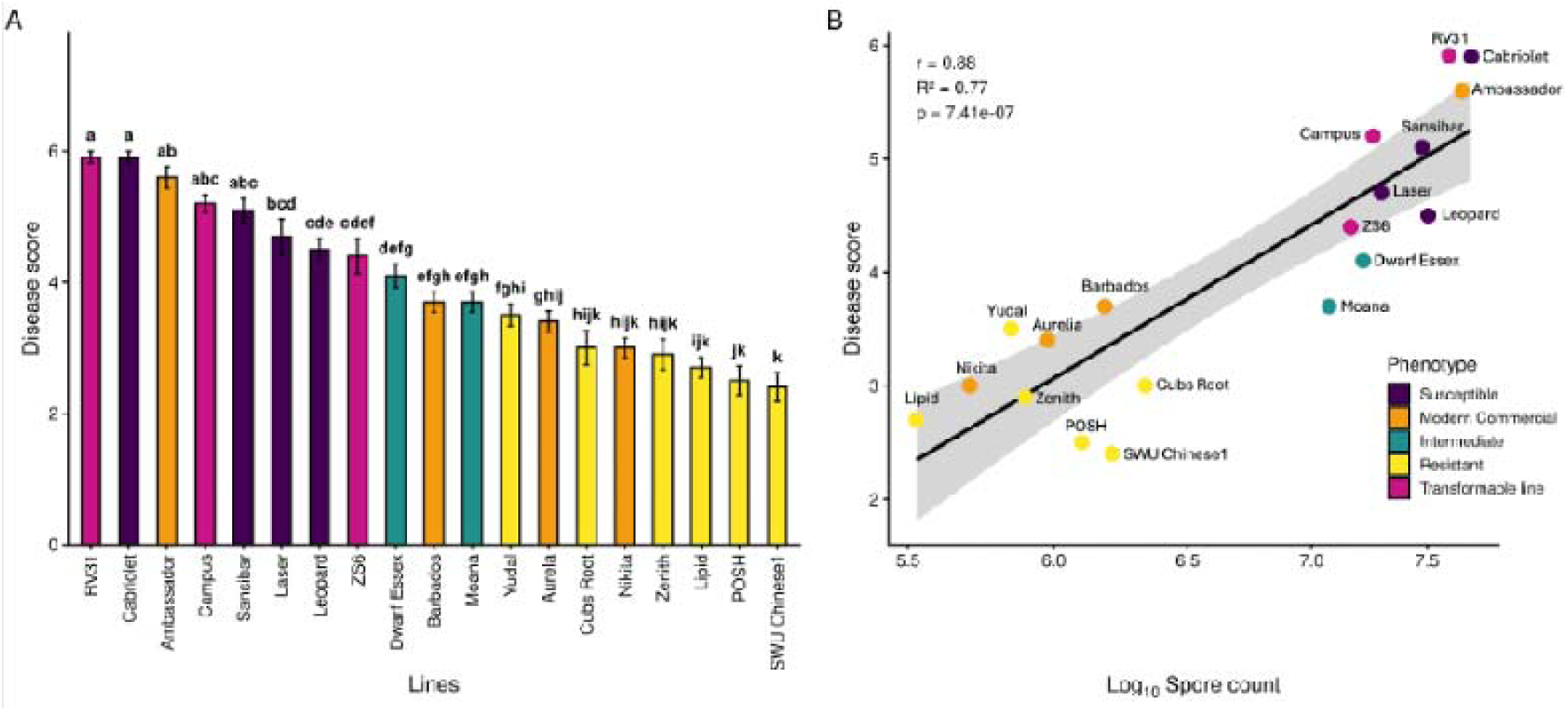
Light leaf spot disease score and its correlation with spore production across *Brassica napus* resistant, intermediate, susceptible, commercial and transformable lines. Seedlings were grown and inoculated under glasshouse conditions. (A) Mean disease scores of *B. napus* cultivars following infection with *Pyrenopeziza brassicae* population collected from Rothamsted, ranked by increasing disease severity. Bars represent mean disease scores ± standard error of the mean (n = 10). Different letters above bars indicate statistically significant differences among cultivars based on analysis of variance followed by Tukey’s HSD test (*P* < 0.05). Cultivars are colour-coded according to phenotype class (purple, green, yellow) and type of accession (orange, pink). (B) Correlation between mean spore counts per 25 cm² leaf area (n = 6) and disease scores across cultivars. Points represent cultivar means, coloured by phenotype class and accession type. The solid line shows the fitted linear regression with the shaded area indicating the 95% confidence interval. Pearson’s correlation coefficient (*r*), coefficient of determination (*R²*), and the associated *P*-value are shown.

Further phenotypic analyses using spore counting (Figure 3A) and quantitative PCR (qPCR) with primers for ITS rDNA of *P. brassicae* (Calderon *et al*., 2002; Figure 3B) confirmed the selected accessions as susceptible (Cabriolet, Sansibar, Laser), intermediate (Moana, Dwarf Essex) or resistant (Cubs Root, POSH, SWU Chinese1). Regression analysis of spore count and qPCR derived pathogen DNA concentration identified a significant positive correlation between phenotyping methodologies, confirming the different resistance classification of these accessions (Figure 3C).

**Figure 3.**
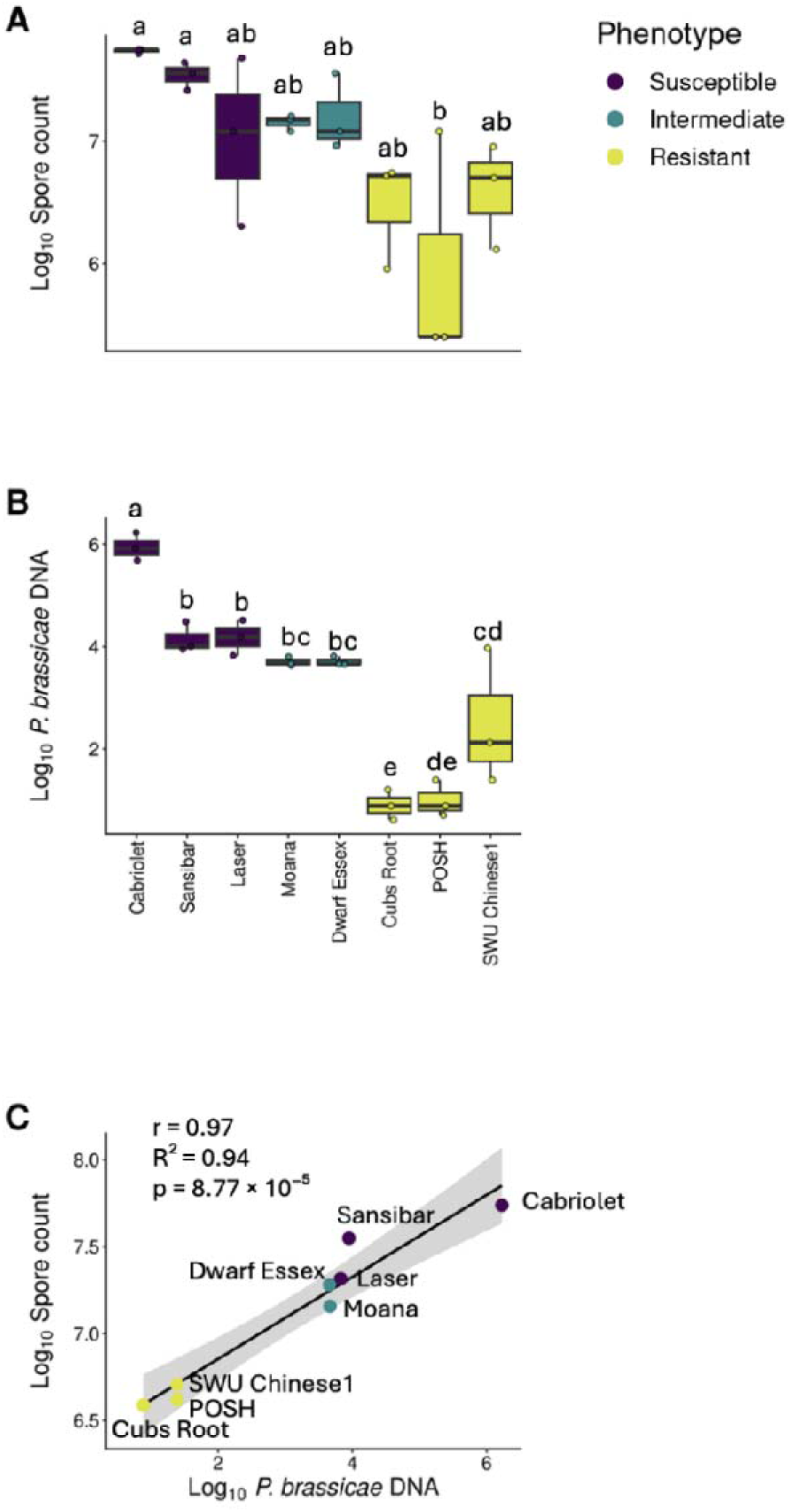
Susceptibility analysis of eight selected *Brassica napus* lines to the light leaf spot pathogen *Pyrenopeziza brassicae* collected from Rothamsted. (A) Disease susceptibility of the selected lines based on fungal spore counts (log_₁₀_-transformed). Resistant, intermediate and susceptible lines are colour-coded. Boxes show the median, lower (25%) and upper (75%) quartiles, with individual points representing biological replicates from two independent glasshouse experiments (n = 6). (B) Disease susceptibility of the selected lines based on the amount of *P. brassicae* DNA (pg) in infected leaves (log_10_-transformed), with susceptibility categories colour-coded. (C) Linear regression analysis of spore counts per 25 cm² leaf area versus *P. brassicae* DNA across the selected lines.

### Resistant *B. napus* line Cubs Root limits *P. brassicae* hyphal growth and colonisation

To examine whether the observed disease resistance affects early infection or subsequent colonisation by *P. brassicae*, pathogen growth and development during foliar colonisation of *B. napus* were monitored on the resistant spring oilseed rape doubled haploid (DH) line, Cubs Root, and the susceptible cultivar, Cabriolet (Figure 4). Using trypan blue staining spores were visible immediately after foliar inoculation (20 min) and at 1 day post-inoculation (dpi), with germination observed at 1 dpi, followed by host cuticle penetration at 2 dpi, hyphal branching at 4 dpi and subcuticular colonisation by 8 dpi. No differences in spore germination or penetration were evident between cultivars during the first two days after inoculation. However, hyphal branching and subcuticular colonisation differed between the two *B. napus* lines at 4 and 8 dpi, with more extensive fungal growth observed in the susceptible cultivar Cabriolet than in the resistant Cubs Root (Figure 4A).

**Figure 4.**
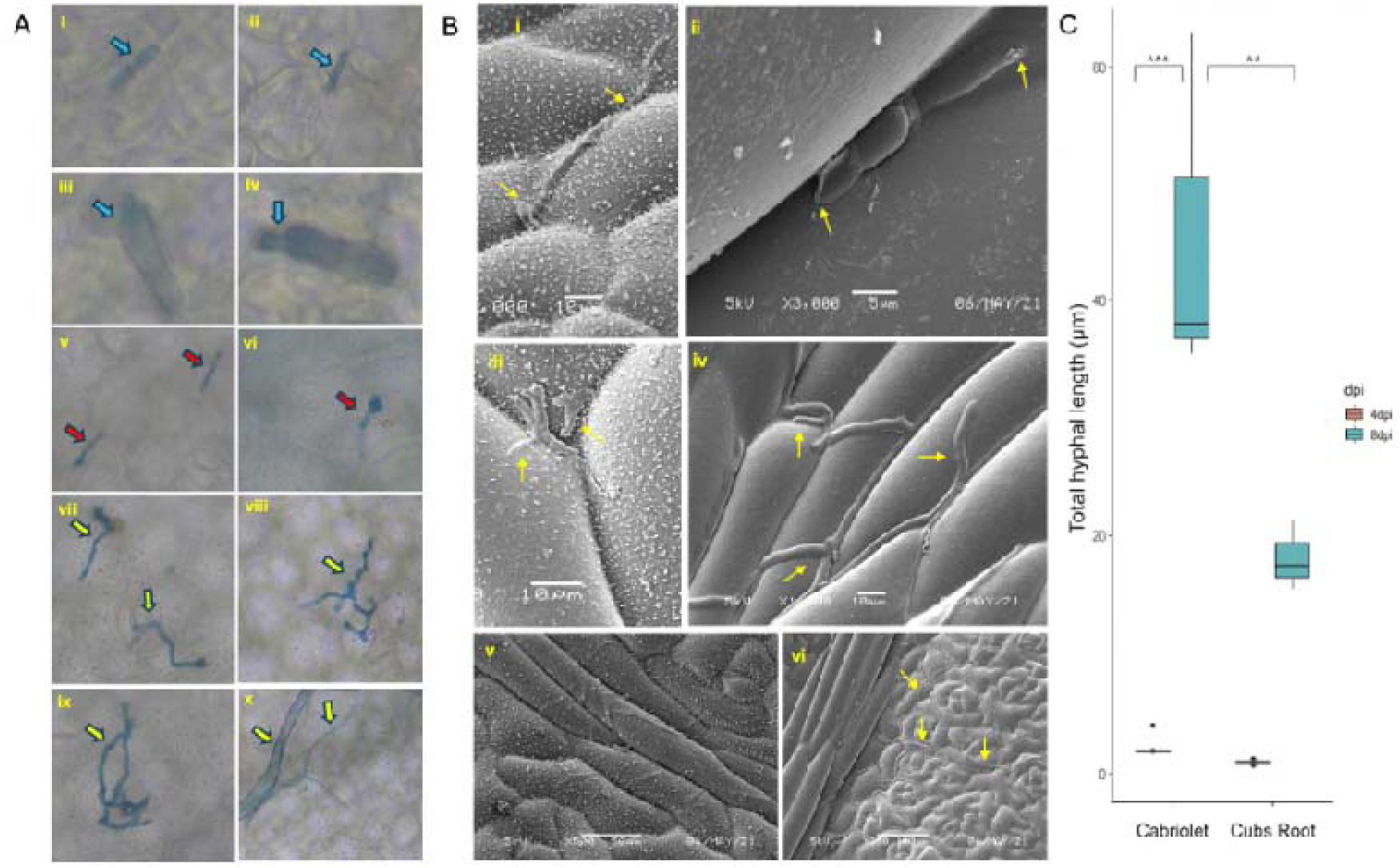
Microscopic analysis of *Pyrenopeziza brassicae* infection in *Brassica napus*. A) Trypan blue stained images showing the progression of *P. brassicae* infection in a resistant line, Cubs Root (i, iii, v, vii, ix), or a susceptible cultivar, Cabriolet (ii, iv, vi,viii, x), at successive time points after inoculation. Arrows indicate *P. brassicae* spores or hyphae. (B) Scanning electron micrographs showing cuticular penetration and subsequent development of *P. brassicae* in the resistant line Cubs Root. Yellow arrows indicate subcuticular hyphae. (C) Quantitative analysis of *P. brassicae* hyphal lengths in infected leaves of *B. napus* at 4 dpi and 8 dpi. The lengths of *P. brassicae* hyphae were measured using SEM images of infected leaves of susceptible cv. Cabriolet or resistant line Cubs Root and ImageJ software. Three micrographs each of 100 μm^2^ were used. A significant difference in total hyphal length between the resistant cv. Cubs Root and susceptible cv. Cabriolet was observed at 8 dpi (*P* < 0.01,**), with no significant difference at 4 dpi. Additionally, within the susceptible cv. Cabriolet, hyphal length differed significantly between 4 dpi and 8 dpi (*P* < 0.001,***).

Scanning electron microscopy analysis agreed with the trypan blue stained microscopic observations, revealing similar early infection processes in both resistant and susceptible lines (Figure 4B, 5B). Again, hyphal branching (4 dpi) and subcuticular colonisation (8 dpi) appeared more extensive in Cabriolet than in Cubs Root. Colonisation of Cabriolet at 8 dpi initially occurred with subcuticular hyphae spreading along veins (Figure 5A) and into the leaf lamina (Figure 4B). Quantification of hyphal length demonstrated significantly increased colonisation relative to resistant line Cubs Root at 8 dpi (*P* < 0.01, Figure 4C). These data illustrate that although germination and penetration did not differ between these *B. napus* lines, resistance responses in Cubs Root reduced the growth and development of the fungal pathogen during hyphal branching and colonisation from 4 dpi onwards.

**Figure 5.**
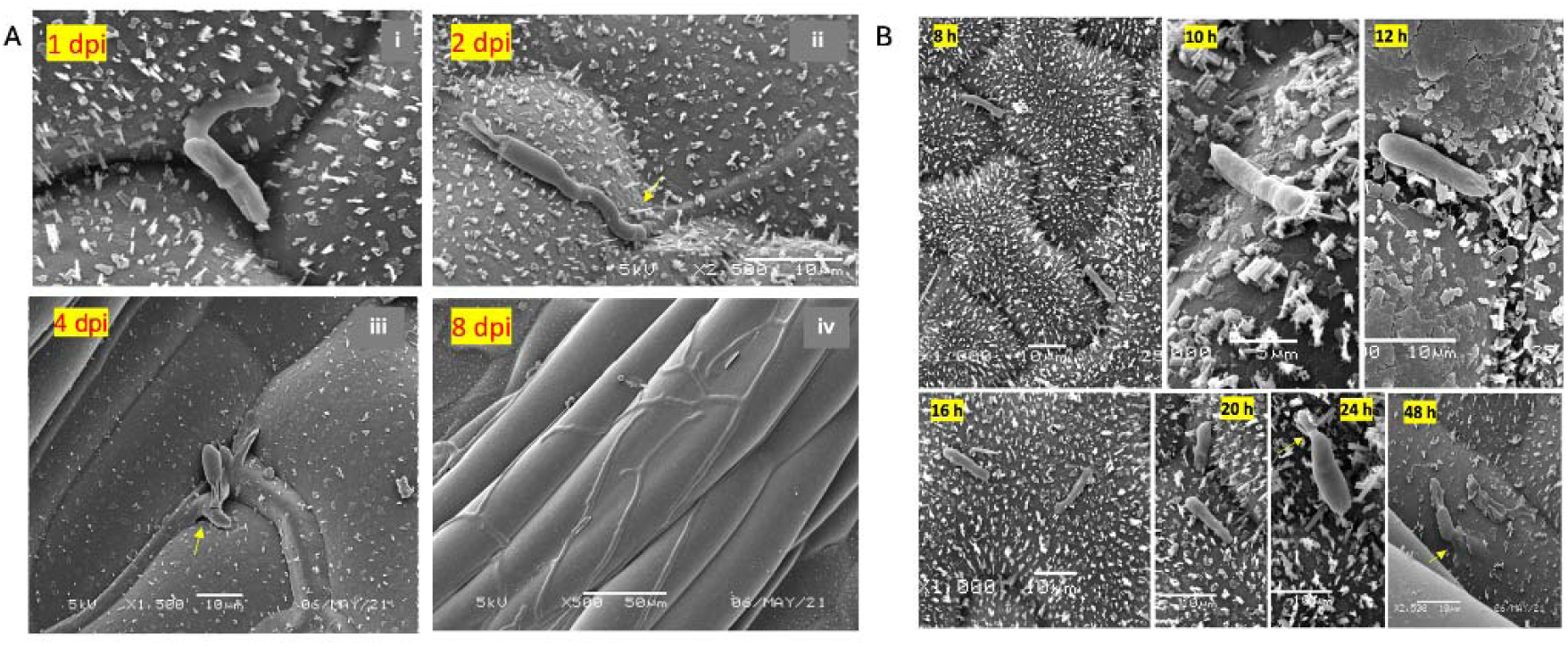
Ultrastructural characterisation of *Pyrenopeziza brassicae* pathogenic development in *Brassica napus*. (A) Scanning electron micrographs (SEMs) illustrating major stages of *P. brassicae* pathogenicity in the susceptible *B. napus* cultivar Cabriolet. Spores germinate at 1 dpi (i), penetrate the cuticle at 2 dpi (ii), initiate hyphal branching at 4 dpi (iii), and establish colonisation by 8 dpi (iv). (B) Scanning electron micrographs showing early interactions between *P. brassicae* and the susceptible cv. Cabriolet. Spores are visible from 8 to 20 h post-inoculation (hpi), begin to germinate at 24 hpi, and penetrate the leaf surface by 48 hpi.

### Pathogen-induced expression of *BAHD acyltransferase* in susceptible *B. napus* lines contrasts the regulation of other gene expression markers

Fell *et al*. (2023) previously identified eight genes where expression level in unchallenged material was associated with quantitative resistance against *P. brassicae*. To determine if differences in expression were present across the four distinct infection stages, six lines spanning the disease spectrum were inoculated with *P. brassicae*: Cabriolet, Sansibar and Laser (susceptible) and Cubs Root, POSH and SWU Chinese1 (resistant). Gene expression was then quantified using multiplex real-time PCR (qRT-PCR). No amplification was detected for the *SNF1 kinase homolog 10 (KIN10)* using TaqMan primers, as confirmed by agarose gel electrophoresis (Supplementary Figure 2).

Differences in gene expression were identified between susceptible and resistant *B. napus* lines following inoculation (Figure 6). *PR1* was more highly induced in resistant cultivars than in susceptible ones, with early activation in Cubs Root, POSH and SWU Chinese1 at 1-2 dpi, perhaps indicative of early salicylic acid (SA) signalling (Haddadi *et al*., 2016). *C4H* expression was induced in the resistant lines and most markedly and persistently in POSH, with transient induction in Cubs Root and SWU Chinese1. Induction of *PLC4* was strong and sustained only in the resistant line POSH. Pathogen-induced expression of *40S ribosomal subunit protein S24* was similar to *C4H*, although SWU Chinese1 was not responsive. The *β-adaptin* gene was the most rapidly and transiently responding gene, with peaks of expression at 1 dpi during spore germination. All resistant lines shared this characteristic, although expression of susceptible cv. Laser was later at 2 dpi and less pronounced. *USP* was the second fastest responding gene and was even induced at 1 dpi in Laser.

**Figure 6.**
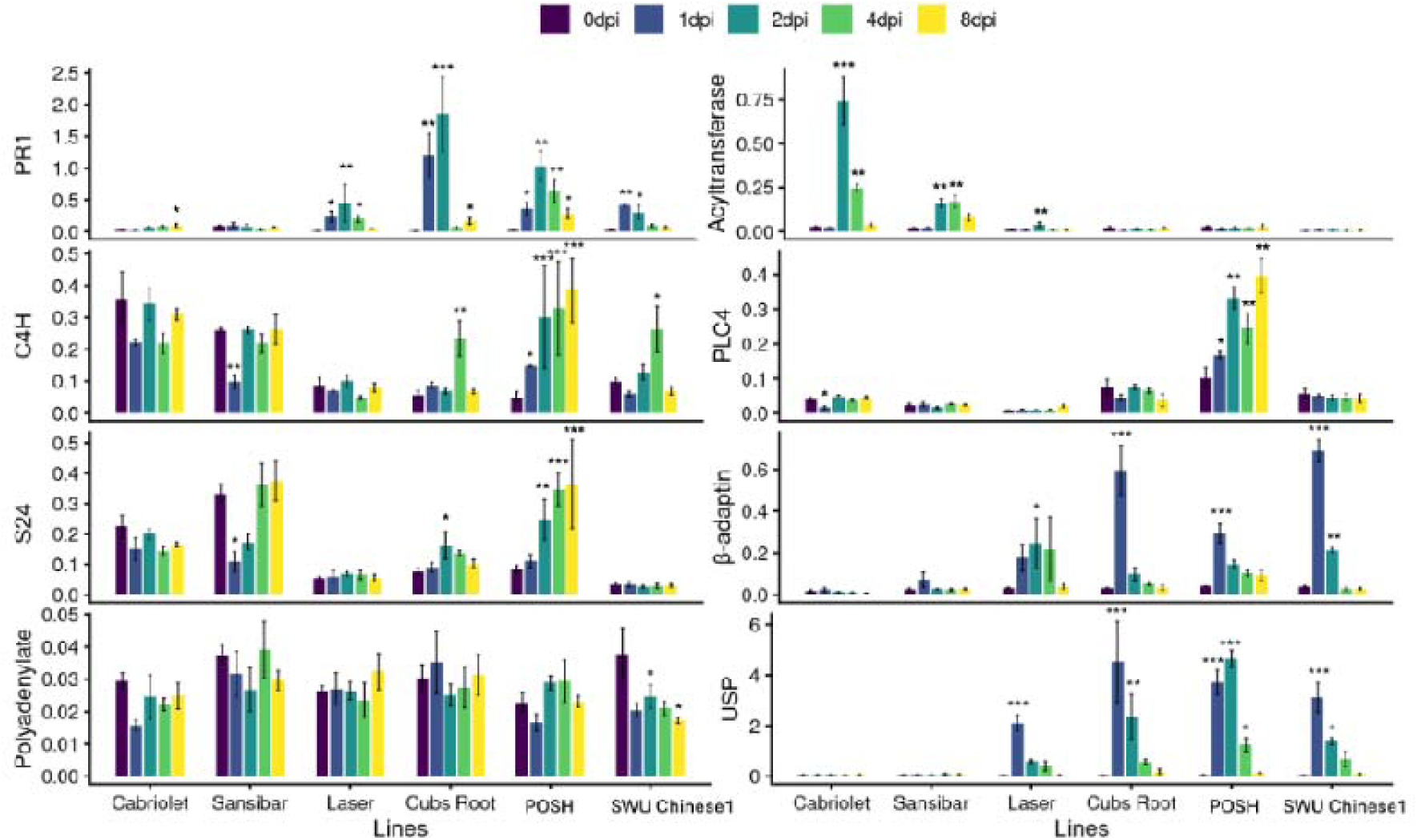
Normalised relative expression of gene expression markers and *PR1* during *Pyrenopeziza brassicae* infection of *Brassica napus* lines. Normalised relative quantity (NRQ) of eight genes: *PR1*, *BAHD acyltransferase*, *cinnamate-4-hydroxylase* (*C4H*), *phosphatidylinositol-specific phospholipase C* (*PLC4*), *40S ribosomal subunit protein S24*, *β-adaptin*, *polyadenylate-binding protein 1*, and *universal stress protein* (*USP*). Bars show means ± standard deviations of three biological replicates, each measured with two technical replicates. Asterisks indicate significant differences from the 0 dpi control within each line based on Dunnett’s test (*P* < 0.05,*; *P* < 0.01,**; *P* < 0.001,***).

Notably, gene expression patterns in the line POSH and cv. Laser were unique. Induced expression of *C4H*, *PLC4* and *40S ribosomal subunit protein S24* was stronger in POSH than all the other lines, suggesting that the defence response might be different. POSH is a resynthesized *B. napus*, developed via induced hybridisation of *B. rapa* and wild *B. oleracea* (Wood, 2010). It may therefore be a valuable source of novel resistance absent from the commercial oilseed rape gene pool. Some induction of *PR1*, *USP* and *β-adaptin* occurred in cv. Laser, although to a lesser degree than in the resistant lines. This concurs with the disease phenotypes, in that cv. Laser was less susceptible than cvs. Cabriolet and Sansibar.

In contrast, *BAHD acyltransferase* expression was significantly induced only in the susceptible cvs. Cabriolet, Sansibar and Laser, without any significant expression in resistant lines. This pattern of expression was different from all other genes analysed. It is therefore suggested that pathogen-induced *BAHD acyltransferase* expression is associated with disease progression. This latter finding reinforces our previous assumption that *BAHD acyltransferase* may encode an enzyme that confers susceptibility to *P. brassicae*.

### Potential contributions of aromatic and aliphatic glucosinolates to defence against *P. brassicae*

GSLs are key defence metabolites in Brassicaceae and play an important role in resistance against bacterial and fungal pathogens (Bednarek *et al*., 2009; Clay *et al*., 2009). To assess the association of GLSs with fungal colonisation, total foliar GSL content was quantified in eight contrasting *B. napus* lines spanning the disease spectrum (Cabriolet, Sansibar, Laser; Moana, Dwarf Essex; Cubs Root, POSH, SWU Chinese1). When comparing seedlings at growth stage 1,6 (Sylvester-Bradley, 1985), total GSL concentrations were observed to vary in leaves of different developmental stages (Supplementary Figure S3). Total GSL concentrations decreased with age; younger leaves (true leaf number 5) had the highest GSL content, indicative of the protective function of these natural plant products in young developing *B. napus* leaves against biotic stress (Badenes-Perez *et al*., 2014).

Total foliar GSL content differed between susceptible and resistant *B. napus* lines (Figure 7A). Leaves of susceptible cvs. Cabriolet, Sansibar and Laser contained the lowest GSL levels, although the GSL content of resistant line SWU Chinese1 was not significantly higher than these susceptible lines. Fodder-type cv. Moana contained intermediate GSL levels, consistent with its intermediate disease susceptibility rankings based on spore counts and pathogen biomass (Figure 3), although GSL content was not significantly different from Cubs Root. The highest GSL amounts in leaves were found in POSH and Dwarf Essex, consistent with the fact that POSH is resistant against *P. brassicae*, although the forage rape Dwarf Essex was ranked intermediate in susceptibility to this fungal pathogen (Figure 7A).

**Figure 7.**
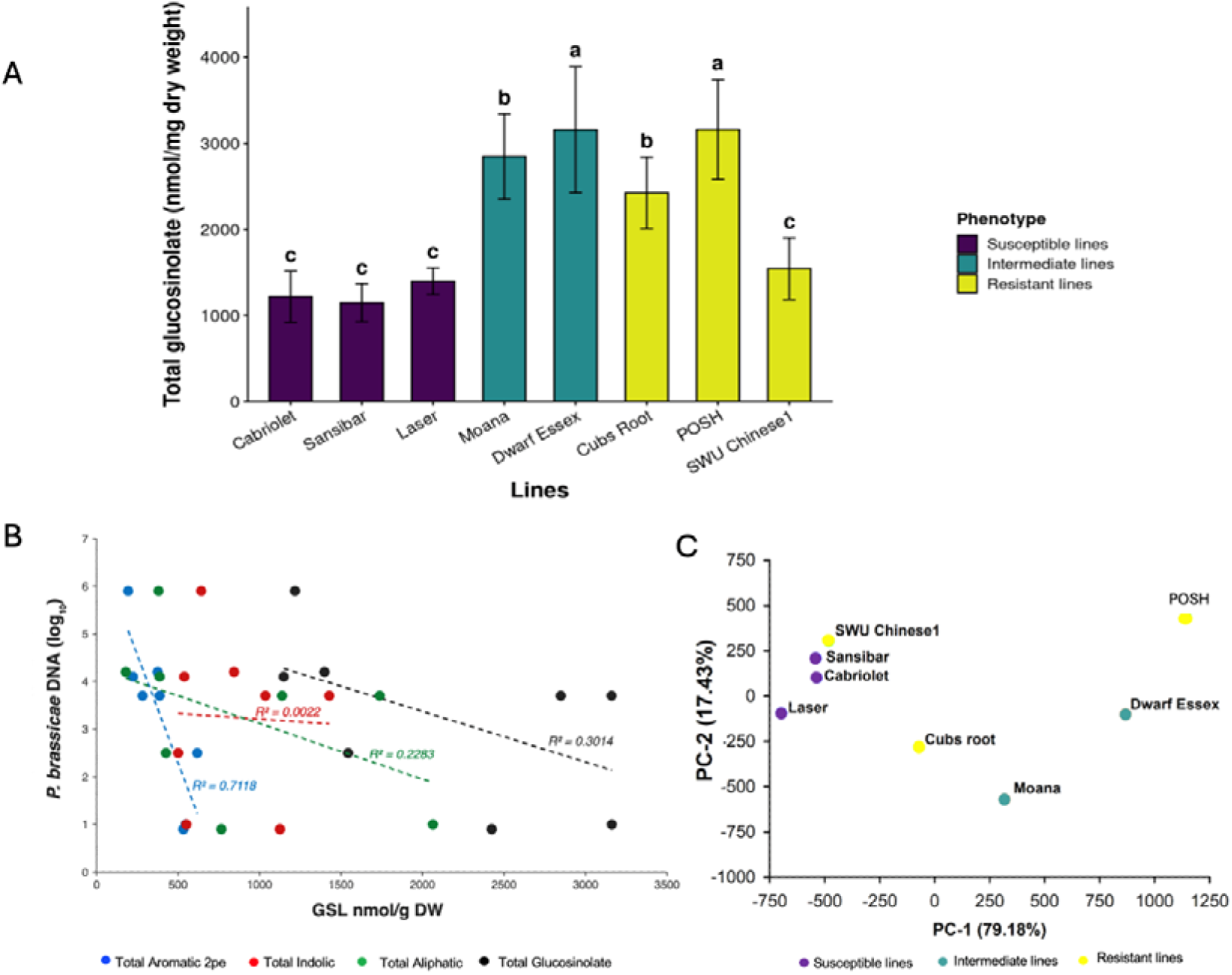
Glucosinolate content in *Brassica napus* lines and relationship with light leaf spot disease severity. (A) Total glucosinolate (GSL) concentrations quantified by HPLC in leaves of *B. napus* lines grouped by disease phenotype (susceptible, intermediate, and resistant). Error bars represent standard errors of five leaves each. (B) Linear regression analysis between total, aromatic 2-phenylethyl (2pe), aliphatic, and indolic GSL concentrations (nmol g⁻¹ dry weight) and pathogen biomass as determined by qPCR of *Pyrenopeziza brassicae* ITS rDNAacross *B. napus* cultivars/lines. Coefficients of determination (*R*²) are indicated. (C) Principal component analysis (PCA) of *B. napus* cultivars/lines based on total aliphatic, aromatic, and indolic GSL profiles. PC-1 (79.18% of variance) and PC-2 (17.43% of variance) are shown. Cultivars/lines are colour-coded according to their susceptibility to the light leaf spot pathogen *P. brassicae*.

These results suggested that although there was a weak correlation between total GSL content and *P. brassicae* biomass in infected leaves (Figure 7B; *P* = 0.096, two-tailed test), with leaves of the susceptible cvs. Cabriolet, Sansibar and Laser containing the lowest GSL amounts.

Further analysis of the different GSL-types revealed that whereas indole GSL content was not correlated with disease scores, negative correlations existed between pathogen biomass and aliphatic (*P* = 0.078, one-tailed test) or aromatic GSL contents in the same eight lines (Figure 7B). In particular, the amounts of 2-phenylethyl GSL (2PE) in leaves were strongly correlated with decreased pathogen biomass (*P* = 0.034, two-tailed test). Principal component analysis (PCA) of GSL-types showed that the first principal component (PC-1) and the second principal component (PC-2) accounted for 79.18% and 17.43% of the variance in GSL content, respectively (Figure 7C). The susceptible cvs. Cabriolet, Sansibar and Laser had the lowest PC-1 scores, i.e. aliphatic GSL levels. Resistant line POSH had the largest PC-1 score, but SWU Chinese 1 had a low PC-1 score, with Cubs Root being intermediate. Nevertheless, SWU Chinese1 had the second highest PC-2 score and contained the most amount of aromatic GSL (Supplementary Table 5). Levels of specific GSL-types in lines Dwarf Essex and Moana were generally intermediate, consistent with their moderate susceptibility phenotypes.

Amongst individual GSLs (Supplementary Table 5), the foliar concentrations of four GSLs were significantly different in specific lines that were tested (Figure 8). 7-Methylsulfinyl heptyl (7MSH) GSL levels were significantly higher in resistant POSH than in susceptible cv. Cabriolet and intermediate Dwarf Essex (Figure 8A). 7MSH GSL concentrations were also negatively correlated with LLS disease severity; this correlation was weakly significant when considering a one-tailed test (*R=-0.548, P=0.080,* Supplementary Figure 4). 4-Methoxy-indolyl-3-methyl (4MO-I3M) GSL was more abundant in POSH than in SWU Chinese 1 (Figure 8B). 5-Methylthiopentyl (5MTP) GSL concentrations were higher in resistant line Cubs Root, intermediate Moana and susceptible cv. Sansibar than in susceptible cv. Laser (Figure 8C). N-methoxy-indolyl-3-methyl (NMO-I3M) GSL concentrations were higher in Cubs Root than in POSH (Figure 8D).

**Figure 8.**
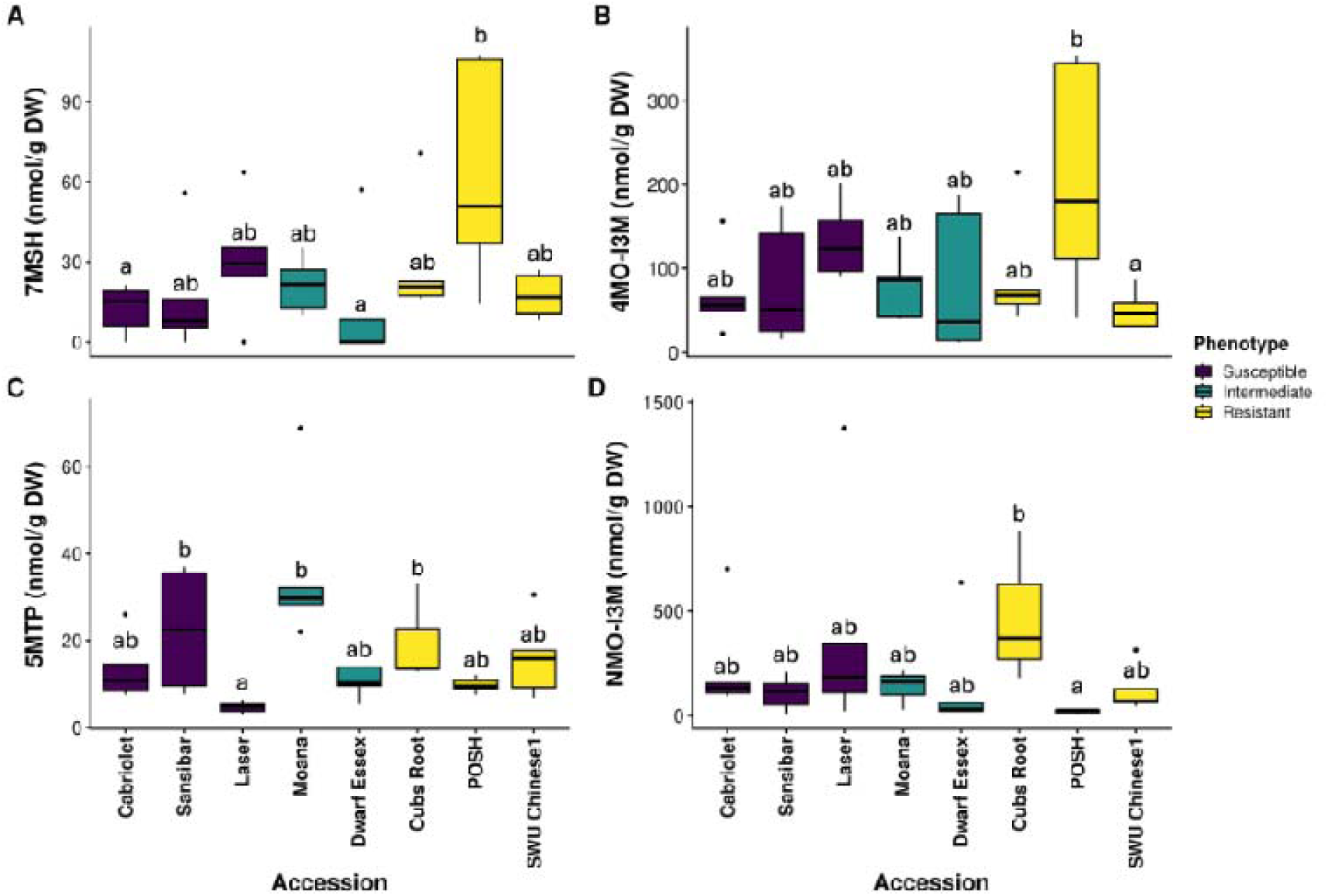
Aliphatic glucosinolate (GSL) concentrations in *Brassica napus* lines with different levels of susceptibility to *Pyrenopeziza brassicae*. (A) 7-Methyl sulfinyl heptyl (7MSH), (B) 4-methoxy-indolyl-3-methyl (4MO-I3M), (C) 5-methyl thiopentyl (5MTP) and (D) N-methoxy-indolyl-3-methyl (NMO-I3M) GSL concentrations were quantified from freeze-dried leaf samples using HPLC. Five biological replicates were used. Values with the same letters are not significantly different after (A,B) analysis of covariance (ANCOVA) and Tukey’s Honestly Significant Difference (HSD) or (C,D) non-parametric Kruskal-Wallis rank sum test and Dunn’s test of multiple comparisons. Boxes show the median, lower (25%) and upper (75%) quartiles; whiskers show values outside the middle 50% and black dots are outliers.

## Discussion

This study provides an integrated phenotypic, molecular and biochemical characterisation of QDR against *P. brassicae* in *B. napus*, combining controlled-environment phenotyping, pathogen quantification, microscopy, GEM profiling and glucosinolate analysis. Together, these datasets reveal that resistance to light leaf spot (LLS) in *B. napus* is mediated by restriction of pathogen proliferation after penetration, underpinned by coordinated transcriptional and metabolic responses.

### Discrepancies exist between field and controlled-environment resistance

This report includes an analysis of LLS disease phenotypes for new commercial cultivars and transformable lines of *B. napus*. Several cultivars exhibited resistance phenotypes under glasshouse conditions that were inconsistent with their UK Recommended List (RL) ratings (AHDB Cereals & Oilseeds, 2016). Amongst the commercial cultivars, Barbados and Aurelia with a Recommended List LLS resistance score of 8 and Nikita with a Recommended List score of 7 were all classified as resistant under glasshouse conditions based on disease scoring and counting of conidia from surface washes of *P. brassicae*-infected leaves. On the contrary, Ambassador, with a Recommended List score of 7, was highly susceptible to *P. brassicae* under both glasshouse and field conditions. Discrepancies between field performance and glasshouse trials are not unexpected and may reflect genotype × environment interactions, differences in pathogen populations, or stage-specific resistance mechanisms, as previously suggested for LLS resistance (Boys e*t al*., 2007; Karandeni Dewage *et al*., 2018a). Collectively, these findings highlight the importance of complementing field-based RL data with controlled inoculation assays when dissecting genetic resistance mechanisms.

Eight lines were selected to better understand LLS disease susceptibility, *P. brassicae* invasion, molecular and biochemical aspects of host susceptibility/defence. Using qPCR-based quantification of pathogen biomass, it was conclusively shown that cvs. Cabriolet, Sansibar and Laser were susceptible whereas Cubs Root, POSH and SWU Chinese1 were resistant, with Moana and Dwarf Essex being intermediate. Moana and Dwarf Essex were previously classified as susceptible and resistant, respectively (Fell et al., 2023).

The transformable lines RV31, Campus and ZS6 were all susceptible to *P. brassicae*, although to different degrees. These characteristics provide a base line for the functional analysis of genes involved in both resistance, via induced or over expression approaches, or susceptibility, using RNAi and genome editing.

### Resistance restricts pathogen development post-penetration

Microscopy revealed that resistance does not prevent spore germination or initial cuticle penetration, which occurred similarly in both resistant and susceptible cultivars. Instead, the resistant line Cubs Root restricted subcuticular colonisation and proliferation, whereas hyphae of *P. brassicae* extensively branched and spread within the subcuticular space of susceptible cv. Cabriolet. These early observations of effects on fungal growth at 4 dpi and 8 dpi precede observations of delayed resistance in cv. Imola (Boys *et al*., 2012).

Quantitative hyphal length measurements further confirmed that significant differences between resistant and susceptible lines emerged only during later stages of colonisation at 8 dpi, reinforcing the concept that QDR against *P. brassicae* acts primarily by limiting pathogen growth rather than blocking infection outright.

### Defence-associated gene expression supports a quantitative resistance model

The expression of seven out of the eight GEMs previously associated with variation in disease score, was differentially regulated in resistant and susceptible *B. napus* lines after *P. brassicae* infection, implicating their role in QDR or disease susceptibility. *PR1* was used as a marker gene for defence, and consistent with this assumption, was induced during fungal penetration primarily in resistant lines.

Pathogen-induced expression of *C4H*, a key enzyme in the lignification pathway, in resistant lines implicates secondary cell wall fortification and lignification in the restriction of pathogen spread, a defence mechanism widely reported in quantitative resistance systems (Becker et al., 2017, El Houari et al., 2021). Pathogen-induced expression of the *40S ribosomal subunit protein S24* gene in resistant lines POSH and Cubs Root could be indicative of translational reprogramming, resulting in the expression of specific defence-related proteins (Fakih *et al*., 2023). *β-Adaptin* was the most rapidly induced GEM after *P. brassicae* infection in resistant *B. napus* lines and peaked at 1 dpi during the protrusion of fungal germ tubes. β-Adaptins are involved in vesicle trafficking which may include defence-related metabolite transport, consistent with emerging evidence that secretory pathways contribute to plant immunity (Ichino & Yazaki, 2022).

In stark contrast, the *BAHD acyltransferase* was only pathogen-induced in susceptible cultivars. This regulated expression increases our knowledge of this putative susceptibility gene that was already suggested to function as such based on the correlated expression with disease severity in a diverse set of 195 *B. napus* accessions (Fell *et al*., 2023). It is hypothesized that this BAHD acyltransferase contributes to a specific metabolic shift that favours pathogen development rather than host defence.

### Relationship between foliar glucosinolates concentrations and light leaf spot disease severity

GSLs act as phytoanticipins in the Brassicaceae for protection against pests and diseases through engagement of the mustard oil bomb (Stotz *et al*., 2011b). Foliar glucosinolate concentrations in *B. napus* lines with different degrees of susceptibility to *P. brassicae* were therefore studied. Having identified a moderately negative correlation between total GSL concentrations and LLS disease, three GSL-categories were examined. Whereas the relationships between total indole and total aliphatic GSL and disease severity were weak and moderate, respectively, a strong inverse correlation between the aromatic 2PE GSL and the LLS score was observed. PCA confirmed these findings with clustering of susceptible, intermediate and resistant lines. Susceptible cultivars were particularly low in PC-1 that explained >79% of the variance, consistent with the lowest total aliphatic glucosinolate levels in cvs. Cabriolet, Sansibar and Laser. The former two cultivars also had the lowest aromatic glucosinolate levels.

Amongst individual GSLs, 7MSH GSL stands out because, like 4MO-I3M GSL, its concentrations were high in POSH and negatively correlated with LLS disease severity. Chain-elongated aliphatic GSL were previously shown to confer resistance against the necrotrophic pathogen *Sclerotinia sclerotiorum*, generating isothiocyanate catabolites that interfere with fungal growth (Stotz *et al*., 2011). Importantly, total indolic GSLs did not correlate with resistance in this study, suggesting that not all GSL classes contribute equally to defence against *P. brassicae*. Amongst individual indole GSLs, however, there were slight differences in correlation with LLS disease severity. Notably, 4MO-I3M was identified as the most interesting indole glucosinolate because of its high levels in POSH and weak negative correlation with LLS severity. 4MO-I3M has previously been linked to innate immunity (Bednarek et al., 2009; Clay et al., 2009) and resistance against insect pests (Pfalz et al., 2009). The specific involvement of aromatic, aliphatic and indole GSLs points to a more targeted role of secondary metabolites in QDR.

### Integration of genetics, transcription and metabolism in QDR

By integrating phenotypic screening, microscopy, gene expression and metabolite profiling, this study provides a systems-level view of QDR to *P. brassicae*. The convergence of transcriptional regulation (e.g. PR1, PLC4, C4H), cellular trafficking (β-adaptin), and metabolic defence (glucosinolates) suggests that resistance is controlled by a coordinated network rather than isolated pathways. These findings are consistent with recent GWAS and associative transcriptomics studies identifying loci linked to both resistance and glucosinolate biosynthesis (Fell et al., 2023), including a cytochrome P450 gene associated with seed GSL content. Together, these results reinforce the hypothesis that metabolic and signalling pathways are genetically intertwined in conferring durable resistance.

### Implications for breeding and future research

Management of LLS in *B. napus* relies heavily on azole fungicides, against which *P. brassicae* populations have already shown reduced sensitivity (Carter *et al*., 2014; Karandeni Dewage *et al*., 2018), raising the prospect of widespread control failure if chemical reliance continues. The identification of defence-associated genes and metabolite profiles linked to resistance provides valuable targets for breeding programmes aiming to enhance durable resistance to LLS. Future work should prioritise functional validation of candidate genes using gene editing or transgenic approaches, as well as exploring the regulatory links between glucosinolate metabolism and defence signalling. Moreover, dissecting how environmental conditions influence resistance mechanisms will be crucial for translating these findings into field-level improvements.

Deploying durable, QDR-based resistance informed by the *in planta* gene expression markers and glucosinolate associations described here offers an alternative route that reduces selection pressure on the pathogen, lowers the input cost and environmental footprint of oilseed rape production, and reinforces integrated pest management approaches. Combining these molecular markers with marker-assisted breeding therefore represents a practical step towards minimising fungicide dependence and delivering more durable, sustainable LLS control in *B. napus*.

### Experimental Procedures

#### Plant material

Twelve accessions were shortlisted for further study (Cabriolet, Cubs Root, Dwarf Essex, Laser, Lipid, Leopard, Moana, POSH, Sansibar, SWU Chinese1, Yudal and Zenith) out of 195 *Brassica napus* accessions previously phenotyped (Fell *et al*., 2023). In addition, four modern commercial cvs. Nikita, Aurelia, Barbados and Ambassador, and the transformable lines ZS6, Campus and RV31 were assessed.

For the field trial conducted at Thriplow, Cambridgeshire (managed by KWS), 21 *B. napus* accessions were selected out of the 195 previously phenotyped (Fell *et al*., 2023). All of the above-mentioned accessions, except for line ZS6, were included in the field trial, with the addition of cultivars Imola, Tapidor and Temple.

#### Pathogen population

The *P. brassicae* population was collected in 2019 from light leaf spot–infected leaves of KWS-grown *B. napus* genotypes, including the reference cultivars Cuillin, Express and Barbados, at a Rothamsted Research field site (51.813125, −0.382005, UK). Only leaves with light leaf spot symptoms were selected. This population was one of two used in our previous study (Fell *et al*., 2023); the first was collected in 2016 from KWS field experiments on the island of Fehmarn, Germany (54.4701, 11.1329) and used for seven glasshouse experiments at Bayfordbury, while the same 2019 Rothamsted population was previously used for three glasshouse experiments at Rothamsted Research. To induce sporulation, leaf samples were covered with moist tissue, sealed in labelled polyethylene bags, and incubated at 4 °C for 3–6 days. Conidia were washed into sterile distilled water, filtered through autoclaved Whatman filter paper, and counted using a Bright-Line haemocytometer under a stereomicroscope (GX Microscopes, XTC-3A1) to determine spore concentration. Inoculum was adjusted to 10^5^ spores/mL and stored at −20 °C in 10mL aliquots, with a subset preserved in 20% sterile glycerol for long-term storage. For subsequent assays, inoculum was produced by spray inoculation of cv. Cabriolet plants, previously grown in a controlled-environment chamber (FITOCLIMA D1200; ARALAB, Portugal) under an 18 h light/6 h dark photoperiod, with 20 °C/18 °C day/night temperatures. For spore production plants were moved to a growth regimen at 12 h light/12 h dark regime at 16/14 °C. Conidia from infected leaves were subsequently harvested to prepare spore suspensions.

The field trial was inoculated with a *P. brassicae* population that originated from Thriplow in 2021.

#### Disease assessment

Seeds were germinated after 2-days stratification (4°C) and seedlings were grown in a glasshouse in a 1:1 mixture of John Innes No. 3 and multipurpose compost (Miracle-Gro; Evergreen Garden Care, Cardiff, UK), first in trays and then in 9 cm pots without additional fertiliser. Conditions were set to 16/14 °C day/night with supplemental LED lighting (200 µmol m^−2^ s^−1^; 12–14.5 h photoperiod), while humidity was monitored but not controlled. Seedlings were inoculated after 4 weeks at growth stage 1,5 (five true leaves). Plants were spray-inoculated with a *P. brassicae* conidial suspension (1 × 10^5^ spores mL^−1^) containing 0.005% Tween 80, applying 1.2 mL per plant. Following inoculation, plants were covered with polyethylene sheets for 48 h to maintain high humidity and incubated under a 12 h light/12 h dark regime at 16/14 °C. At 21 dpi, infected leaves were cold-incubated (4 °C for 5 days) to induce sporulation.

Disease severity was scored using a 1–6 scale (1 = no sporulation, resistant; 2 = <10% leaf area with sporulation, 3 = 10–25% leaf area with sporulation, 4 = 25–50% leaf area with sporulation), 5 = 50–75% leaf area with sporulation, 6 = 75–100% leaf area with sporulation) (Fell *et al*., 2023). Sporulation was determined using three biological replicates per line. Conidia were collected from a standardised leaf area of 25 cm² per sample by washing with sterile water, filtered through Miracloth, and quantified using a haemocytometer.

Fungal DNA quantification was performed to determine fungal biomass in infected leaf tissues. Genomic DNA was extracted from 20 mg freeze-dried leaf tissue using the DNAMITE Plant Kit (Microzone Ltd., UK) with mechanical lysis via metal beads and FastPrep-24 (MP Biomedicals, UK) homogenisation. DNA was eluted in 40–80 µL sterile nuclease-free water, quantified, diluted to 50 ng µL^−1^, and stored at −20 °C. Pathogen DNA was quantified by qPCR using ITS primers (Karolewski *et al*., 2006): PbITS (F) 5′-ttg aac ctc tcg aag aag ttc agt ct-3′ and PbITS (R) 5′-aga ttt ggg ggt tgt tgg cta a-3′. Reactions included two technical replicates per sample, non-template controls (NTC), five DNA standards (1–10,000 pg), and a positive amplification control containing purified *P. brassicae* DNA. qPCR cycling conditions were 95 °C for 2 min; 50 cycles of 95 °C for 15 s, 58 °C for 45 s, and 72 °C for 45 s with data acquisition at 84 °C for 15 s; followed by a final extension of 95 °C for 1 min, 58 °C for 30 s, and 95 °C for 30 s. Ct values were obtained using consistent default thresholds. Technical replicates were averaged when Ct values differed by ≤0.5 cycles; if the difference exceeded 0.5 cycles, the sample was re-analysed. A standard curve (1–10^5^ pg *P. brassicae* DNA) was generated by linear regression of Ct versus log_10_ DNA quantity, and sample pathogen DNA was interpolated and expressed as pg per reaction, then normalised to pg per mg dry leaf tissue.

Field trial experimental samples were collected in March, May and June of 2022, by harvesting 10 leaves per block picked at random, and assessed using both visual assessment and spore counting.

#### Microscopic imaging of fungal infection and growth

Susceptible *B. napus* line Cabriolet and resistant line Cubs Root were analysed to assess fungal structures and host responses. Plants were grown under controlled environment conditions using three biological replicates and spot-inoculated by placing sterile Whatman filter paper squares (0.8 × 0.8 mm) soaked in a conidial suspension (1 × 10^5^ spores mL^−1^) onto marked areas of the third or fourth true leaves. Inoculated leaves were covered with polyethylene sheets to maintain high humidity. Leaf discs were collected at multiple time points post inoculation (20 min, hourly up to 15 h, and daily up to 8 dpi). Trypan blue staining was done using a protocol adapted from established methods for visualising fungal structures in plant tissues (Van Wees, 2008). Stained samples were examined using a GXM-L2800 microscope (GT Vision, UK), and micrographs were captured to assess *P. brassicae* penetration and endophytic growth.

For low-temperature scanning electron microscopy (SEM), samples were prepared using a 2100 Gatan Alto cryo preparation system following the manufacturer’s protocol (Gatan, UK). From spot-inoculated areas, leaf discs (0.5–1 cm²) were excised using a sterile blade and mounted onto SEM stubs using a 1:1 mixture of colloidal graphite (TAAB) and Tissue-Tek (Sakura). Samples were rapidly frozen in liquid nitrogen, transferred into the preparation chamber via a vacuum transfer device, fixed at −90 °C for 2 min and sputter-coated with gold for 1 min. Samples were imaged using a JEOL 6360 low-vacuum scanning electron microscope at an accelerating voltage of 10 kV, with a spot size of 38 and a working distance of 15 mm. Signals were detected using a secondary electron detector, and digital micrographs were acquired using the JEOL SEM Control User Interface (version 6.04).

Spot-inoculated *B. napus* susceptible line Cabriolet was used to assess early host–pathogen interactions under controlled-environment conditions. Samples were collected at 8, 10, 12, 16, 20, 24 and 48 h post-inoculation, with time points selected based on Trypan blue staining results. To investigate differential interactions between susceptible and resistant lines, SEM imaging was also performed at 2, 4, 6 and 8 days post-inoculation using Cabriolet and Cubs Root. Total hyphal length was quantified using ImageJ software to assess differences in colonisation between lines at 4 and 8 dpi. Three micrographs per line (Cabriolet or Cubs Root), each with a scale bar of 100 µm, were used for the measurements. Micrographs were obtained from three independent biological replicates per line.

#### Gene Expression Profiling of Gene Expression Markers (GEMs)

Gene expression profiling of candidate gene expression markers (GEMs) was performed after identifying the *B. napus* gene models of previously detected GEMs in *B. napus* (Fell *et al*., 2023). Orthologous genes were identified by sequence alignment and phylogenetic comparison using published Brassicaceae genome resources (Chalhoub *et al*., 2014). Candidate loci were selected from homoeologous gene lists and confirmed by gene tree analysis to ensure correct orthologue assignment. These alignments were used for homologue specific TaqMan assay design (Supplementary Table S3), which lists assay names, gene targets, and amplicon context sequences. These assays were validated prior to expression analysis by assessing amplification efficiency and linearity from standard curves, melt-curve specificity, and agarose gel electrophoresis of PCR products. Singleplex and multiplex reactions produced comparable Ct values, confirming assay compatibility.

Six *B. napus* lines (Cabriolet, Sansibar, Laser, Cubs Root, POSH and SWU Chinese1) were grown under controlled-environment conditions and spot-inoculated with a *P. brassicae* population. Leaf discs were collected at 1, 2, 4 and 8 days post inoculation (dpi), with mock-inoculated plants serving as controls. Samples were snap-frozen in liquid nitrogen, freeze-dried, and homogenised to a fine powder for 1 min at 27.5 Hz using a FastPrep homogeniser (MP Biomedicals, UK) with three sterile 3 mm metal beads. Total RNA was extracted using the E.Z.N.A.® Plant RNA Kit (Omega Bio-tek, UK) according to the manufacturer’s protocol and treated with RNase-free DNase (Omega Bio-tek, UK) to remove residual genomic DNA. RNA concentration was determined spectrophotometrically (NanoDrop, Thermo Fisher Scientific, UK), and first-strand cDNA was synthesised using the qPCRBIO cDNA Synthesis Kit (PCR Biosystems LTD, London, UK) following the supplier’s instructions.

Gene expression was quantified using multiplex TaqMan qPCR on an ABI 7700 system. Actin was used as the reference gene, and normalised relative quantities (NRQ) were calculated according to Rieu *et al*. (2009).

#### Quantification of glucosinolates

Eight *B. napus* cultivars/lines were grown in a controlled-environment cabinet (FITOCLIMA D1200; ARALAB, Portugal) under a 12 h light/12 h dark photoperiod at 20 °C (day) and 18 °C (night) in a randomised block design with eight biological replicates (plants) per cultivar. Leaves were collected from 30-day-old plants at the 1,5 or 1, 6 true leaf stage (Sylvester-Bradley *et al*., 1984). For each plant, leaves 1–5 (numbered from oldest to youngest) were sampled, with leaf 1 representing the oldest and leaf 5 the youngest true leaf. Leaf samples were freeze-dried for three days prior to glucosinolate analysis.

GSLs were extracted and analysed as desulfo-glucosinolates (dsGSLs) as previously described (Kliebenstein et al., 2001). Briefly, freeze-dried leaf material was extracted with 85% methanol containing pOHB glucosinolate as internal standard and loaded onto DEAE Sephadex. DsGLS were released by adding sulfatase and eluted with milliQ grade water before separation and quantification using UHPLC/TQ-MS (Jensen et al., 2015). Total GSLs or aliphatic, indolic or aromatic subgroups were calculated by adding all individual GSLs or GSLs of each subgroup.

#### Statistical analysis

Statistical analyses were performed using R software version 4.4.2. Data were log10-transformed where appropriate to meet assumptions of normality and homogeneity of variance. For data that violated normality or homogeneity assumptions, non-parametric Kruskal-Wallis tests were applied. For normally distributed data with homogeneous variance, analysis of variance (ANOVA) was performed, followed by Tukey’s HSD post-hoc test for pairwise comparisons when ANOVA P-values were less than 0.05.

For gene expression data, analysis of covariance (ANCOVA) was used to account for the effect of the reference gene, Actin. To assess differences across time points within each genotype, Dunnett’s test was applied, comparing each time point to the 0 dpi control.

ANCOVA was used in R to determine the effect of genotype on individual glucosinolates with leaf position as a covariate using the expression: aov(target GSL ∼ Genotype + Leaf). The R function cor.test was used to determine significance of correlation using the Pearson correlation coefficient and a one-tailed test, assuming that GSL concentrations were negatively correlated with disease score.

## Acknowledgements

The authors would like to thank Dr Chinthani Shanika Karandeni Dewage and Heather Fell for their guidance and advice throughout this work. We are grateful to Professor Bruce D. L. Fitt for his valuable advice and support. We thank KWS for granting access to the trial sites at Rothamsted, and the Rothamsted Research and Bayfordbury glasshouse teams for their technical assistance. We also acknowledge Rothamsted Research for their support with SEM imaging.

## Author contributions

A.M.A.: Conceptualization, investigation, formal analysis, writing – original draft. L.G.M.: Investigation, formal analysis. A.Q.: Conceptualization, formal analysis. C.C. and B.A.H.: Investigation, resources. H.U.S. and R.W.: Conceptualization, funding acquisition, supervision, writing – review & editing. All authors contributed to the final manuscript.

## Data availability statement

The data that support the findings of this study are openly available in the text or in the supporting information. Further information is available from the corresponding author on request.

## Funding

This work was supported by ERA-CAPS/BBSRC grants BB/N005112/1 and BB/N005007/1 (MAQBAT; H.U.S., R.W., B.A.H. and C.C.), BBSRC grant BB/V01725X/1 (H.U.S. and R.W.), and BBSRC Institute Strategic Programme Advancing Plant Health BBS/E/JI/230001A (R.W.). A.M.A. was funded by the Hertfordshire Science Partnership, part-financed by the Hertfordshire Local Enterprise Partnership’s Growth Deal 2, the Chadacre Agricultural Trust, and BSPP Covid Funding. L.G.M. was supported by The Perry Foundation and the University of Hertfordshire. A.Q. was funded by the Quality Research (QR) fund of the University of Hertfordshire and the Newton Fund Impact Scheme, managed by the UK British Council and the Egyptian Science, Technology and Innovation Funding Authority (STDF; Project ID: 43937).

The authors declare no conflict of interest.

## Supporting Information legends

**Figure S1 | Susceptibility of *Brassica napus* lines to *Pyrenopeziza brassicae* under field conditions.** (A) Disease score was ranked in 19 *B. napus* lines when grown in a field plot at Thriplow, Cambridgeshire, UK. Disease severity was scored on a scale of 1 to 6 (Karandeni Dewage *et al.,* 2021) after inoculation with a *P. brassicae* population from the same field in Thriplow collected in the previous year. Means and standard errors are shown (*n* = 4). (B) Foliar spore concentrations were determined. Means and standard errors are shown (*n* = 4). (C) Linear regression analysis of spore counts versus disease scores. Individual points represent mean values for both estimates. The line of best fit is shown and the yellow area represents the 95% confidence interval. Pearson’s correlation coefficient (*R*) and the significance of the correlation (*P*-value) are indicated.

**Figure S2 | Agarose gel electrophoresis of quantitative real-time RT-PCR products generated using TaqMan multiplex primers/assays.** Gene names included *cinnamate-4-hydroxylase* (*C4H*), *phospholipase C* (*PLC4*) and 40S ribosomal subunit protein S24e and *universal stress protein* (*USP*) and *PR1*. *BAHD acyltransferase* was included, as was a non-template negative control. Note that *SNF1 kinase homolog 10* (*KIN10*) and *glyceraldehyde phosphate dehydrogenase* (*GAPDH*) did not yield products. *Polyadenylate-binding protein 1* and *cutinase* generated two bands.

**Figure S3 | Glucosinolate content across leaf developmental stages and in *Brassica* lines.** (A) Glucosinolate (GSL) concentration (nmol per g dry weight) measured across leaf developmental stages in *B. napus*, where leaf 1 represents the oldest and leaf 5 the youngest. Values represent means ± SE across eight cultivars, based on five leaves sampled from the same plant at each developmental stage, with error bars indicating standard errors. Statistical significance was determined by one-way ANOVA followed by Tukey’s HSD post-hoc test; different letters indicate significant differences between leaf stages (*p* < 0.05). (B) Total glucosinolate (GLS) concentration (nmol per g dry weight) measured across five leaf developmental stages in eight *Brassica napus* cultivars (Cabriolet, Sansibar, Moana, Laser, SWU Chinese1, Cubs Root, POSH, and Dwarf Essex), grouped by disease phenotype. Lines were classified as susceptible (purple, n = 4), intermediate (teal, n = 2), or resistant (yellow-green, n = 4) based on their disease scores following inoculation with *P. brassicae*. Leaf 1 represents the oldest and leaf 5 the youngest leaf.

**Figure S4 | Relationship between glucosinolate concentrations and light leaf spot disease severity in *Brassica napus*.** Linear regression analysis between (**A**) 7-methylsulfinyl-heptyl (7msh), (**B**) 4-methoxy-indolyl-3-methyl (4moi3m), (**C**) 5-methylthiopentyl (5mtp) or (**D**) N-methoxy-indolyl-3-methyl (nmoi3m) glucosinolate (GSL) concentrations (nmol g^−1^ dry weight) and light leaf spot (LLS) disease score (scale 1–6) across *B. napus* cultivars/lines. Coefficients of determination (R²) and regression equations are indicated.

**Table S1.** *Brassica napus* cultivars and lines used in this study, their disease phenotype classification, source, and the experiments in which they were included.

**Table S2.** *Brassica napus* homologue genes identified from *B. oleracea* and *B. rapa* orthologues, with corresponding gene IDs and functional descriptions.

**Table S3.** TaqMan multiplex qPCR assay details including vendor assay names, gene names, gene descriptions, amplicon context sequences, assay numbers, and fluorescent dyes used.

**Table S4.** Principal component analysis (PCA) loadings of glucosinolate compound groups for principal components 1 and 2 (PC-1 and PC-2).

**Table S5.** Mean concentrations (nmol g^−1^ dry weight) of individual glucosinolates measured across five leaf developmental stages (leaf 1 to leaf 5) in eight *Brassica napus* cultivars.

## Notes

### Competing Interest Statement

The authors have declared no competing interest.

### Summary of Updates

To correct author name and ORCID

