## Supplementary figures and images for "Characterisation of gene expression markers and glucosinolates during discrete infection stages of *Pyrenopeziza brassicae* in *Brassica napus*"

### Supplemental Figure 1.

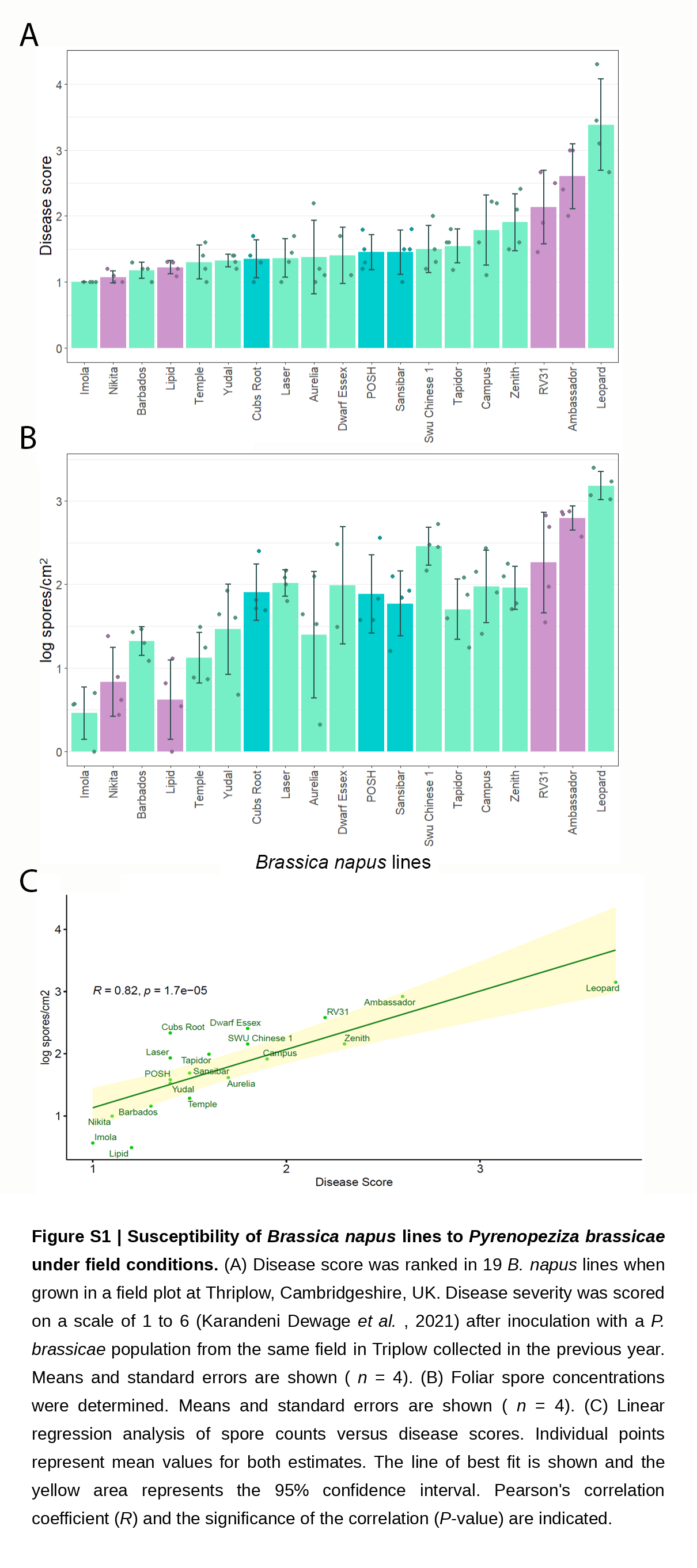

### Supplemental Figure 2.

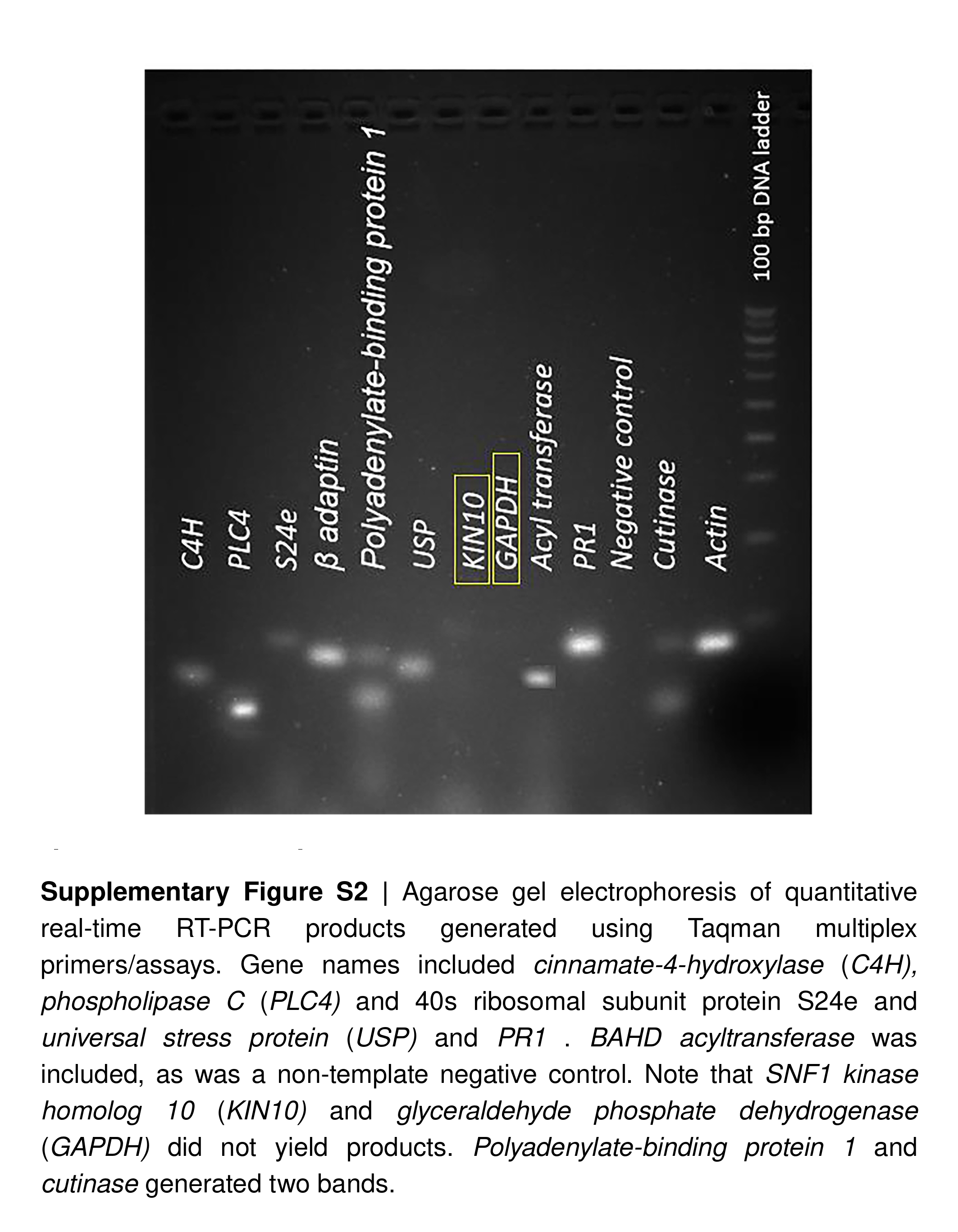

### Supplemental Figure 3.

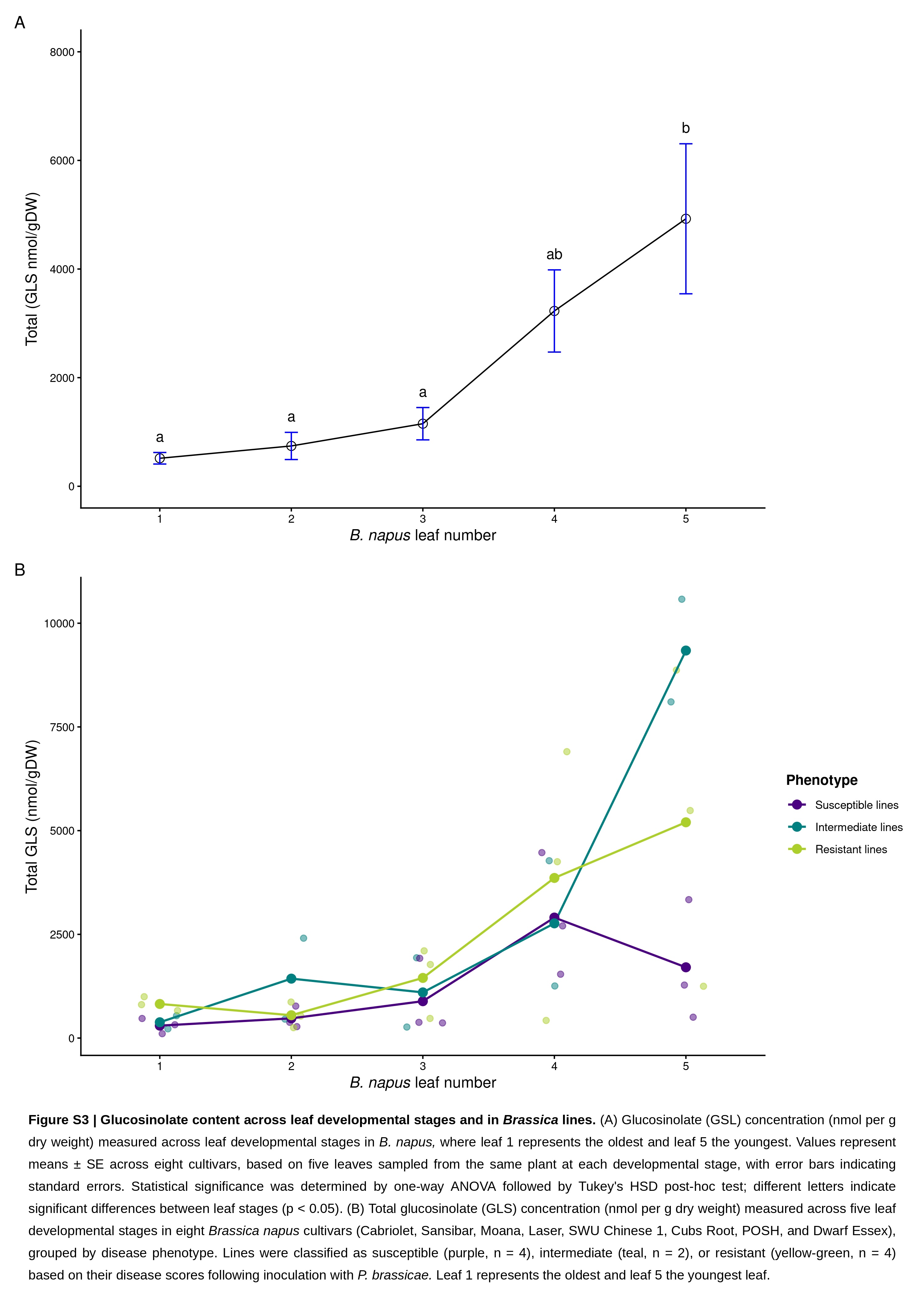

### Supplemental Figure 4.

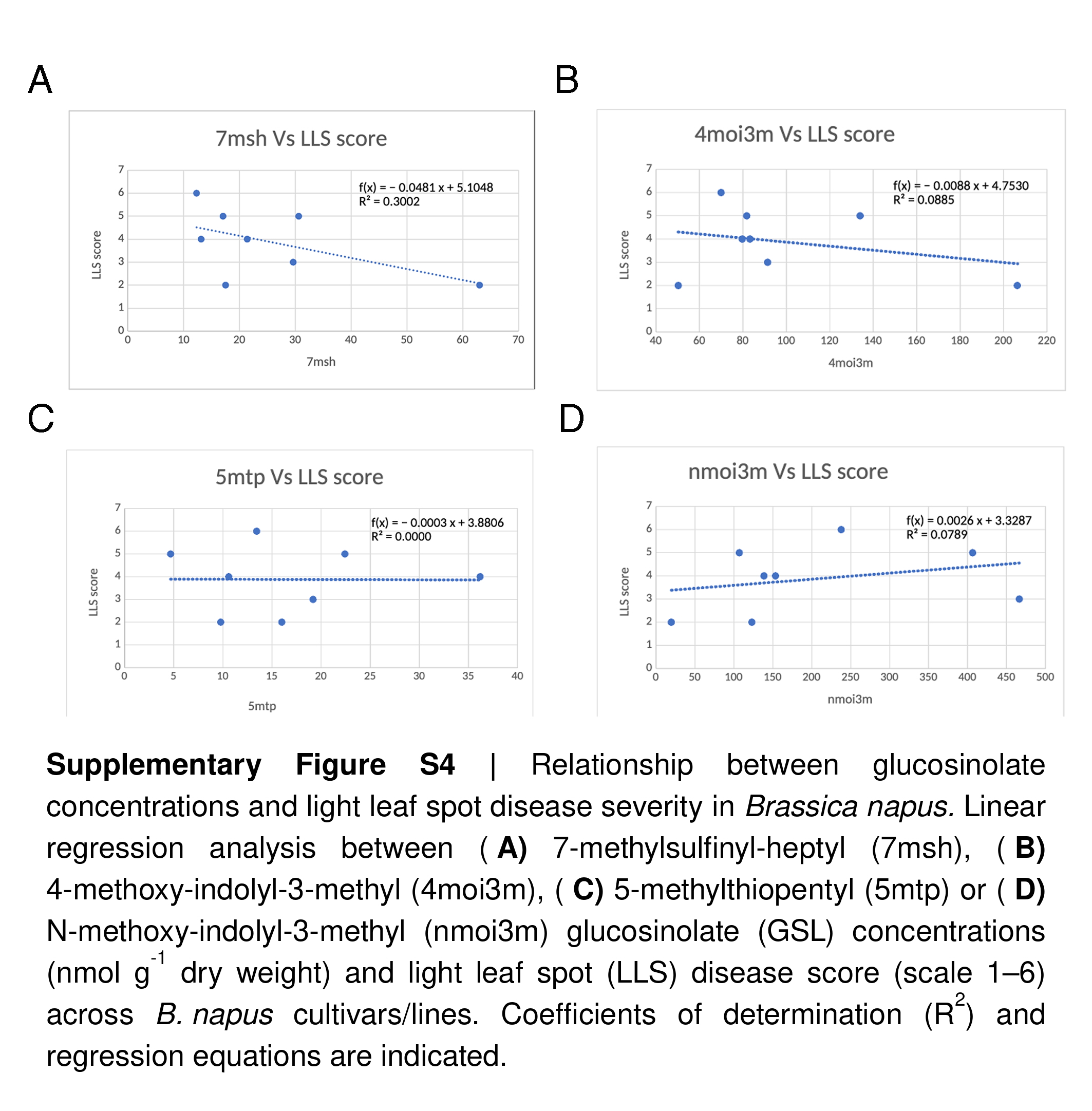
